# Dynamic BMP signaling regulates sclerotome induction and lineage diversification in zebrafish

**DOI:** 10.1101/2024.09.24.614810

**Authors:** Linjun Xie, Roger C. Ma, Katrinka M. Kocha, Emilio E. Méndez-Olivos, Peng Huang

## Abstract

The sclerotome is an embryonic structure that gives rise to various supportive tissues, including the axial skeleton and connective tissues. Despite its significance, the mechanisms underlying sclerotome induction and diversification during embryonic development remain poorly understood. Sclerotome progenitors exhibit transient *bmp4* expression and an active response to BMP signaling. Using BMP gain- and loss-of-function tools, we demonstrate that BMP signaling is both necessary and sufficient for sclerotome induction. Furthermore, through mosaic expression of a dominant-negative tool, we show that BMP signaling induces sclerotome fate in a cell-autonomous manner. Interestingly, different populations of sclerotome-derived cells have distinct BMP signaling requirements. Sclerotome-derived notochord-associated cells in the trunk lack BMP response, and sustained BMP signaling inhibits their differentiation into tenocytes. By contrast, sclerotome-derived fin mesenchymal cells in the fin fold require high levels of BMP signaling for proper morphogenesis. Our findings suggest that dynamic regulation of BMP signaling is crucial for the induction of the sclerotome and the subsequent diversification of sclerotome-derived lineages in zebrafish.

## INTRODUCTION

In vertebrates, the somite is the transient embryonic structure that contributes to diverse cell types in the trunk (Yusuf and Brand-Saberi, 2006; Christ and Scaal, 2009). As it matures, the somite subdivides into the dermomyotome and sclerotome. The dermomyotome further splits into the dermatome and myotome. The dermatome, myotome, and sclerotome give rise to the dermis, muscles, and axial skeleton, respectively. Although the myotome lineage has been extensively studied, the development and diversification of the sclerotome lineage remain poorly understood.

Bone morphogenetic protein (BMP) signaling plays critical roles in many processes during vertebrate embryonic development, including dorsal-ventral patterning, mesoderm formation, and neural patterning (Liu and Niswander, 2005; Dutko and Mullins, 2011; Wang et al., 2014; Yan and Wang, 2021). Secreted BMP ligands bind to transmembrane receptor complexes composed of type I and type II serine/threonine kinase receptors. Upon ligand binding, the constitutively active type II receptor phosphorylates and activates type I receptor, which subsequently phosphorylates Smad1/5/9 proteins. These phosphorylated Smad proteins then translocate into the nucleus to activate target gene expression. BMP signaling is also antagonized by extracellular inhibitors such as Noggin and Gremlin, which sequester BMP ligands and prevent receptor binding (Walsh et al., 2010). During somite development in chick, BMP4 from the lateral plate mesoderm, together with Noggin from the notochord, specifies the somite along the medial-lateral axis (Pourquié et al., 1995; Pourquié et al., 1996; Hirsinger et al., 1997; Tonegawa et al., 1997; Tonegawa and Takahashi, 1998). Active BMP signaling is required for the maintenance of lateral somite, marked by *Pax3* and *cSim1* expression, whereas repression of BMP signaling allows the progression of the myogenic program in the medial somite. Interestingly, treatment of mouse paraxial mesoderm explants with BMP ligands blocks the expression of *Pax1*, a key sclerotome marker (McMahon et al., 1998). Similarly, mouse *Noggin; Gremlin1* double mutants display a loss of *Pax1* expression (Stafford et al., 2011), suggesting that BMP inhibition is required for sclerotome induction. Whether a similar mode of regulation is required for sclerotome development in other species is currently unclear.

In zebrafish, BMP signaling has also been implicated in somite development. Overactivation of BMP signaling delays the myogenic differentiation of dermomyotome cells (Patterson et al., 2010). Different levels of BMP signaling, together with Hedgehog (Hh) signaling, also contribute to the specification of adaxial cells along the dorsoventral axis, resulting in the formation of muscle pioneers (low BMP, high Hh) and superficial slow-twitch muscle fibers (high BMP, low Hh) (Dolez et al., 2011; Maurya et al., 2011; Nguyen-Chi et al., 2012). Interestingly, in addition to adaxial cells, active BMP response has been observed in the dorsal and ventral regions of the somite (Dolez et al., 2011; Maurya et al., 2011). This raises the question of whether BMP signaling plays roles in other somitic compartments.

Our previous work has shown that the sclerotome in zebrafish is characterized by a bipartite organization in the dorsal and ventral regions of each somite (Ma et al., 2018). Interestingly, although Hh signaling is required for the migration and maintenance of sclerotome-derived notochord-associated cells, inhibition of Hh signaling does not impact the induction of the sclerotome domains (Ma et al., 2018). Here, we identify BMP signaling as the key pathway in sclerotome induction. BMP ligands are expressed in or near sclerotome domains, resulting in an active BMP response in sclerotome progenitors. Gain- and loss-of-function manipulations reveal that BMP signaling is both necessary and sufficient for sclerotome induction. Furthermore, BMP signaling is differentially required in distinct sclerotome-derived cell types. Together, our results suggest that dynamic regulation of BMP signaling is essential for the induction and diversification of the sclerotome lineage in zebrafish.

## RESULTS

### BMP ligands are expressed either within or near the sclerotome and its descendants

To investigate the role of BMP signaling in sclerotome development, we examined the expression pattern of several BMP ligands using in situ hybridization (Fig. 1). At 20 hpf, *bmp2b*, *bmp4*, and *bmp6* showed prominent expression along the edge of the fin fold (Fig. 1A). Notably, *bmp4* was also expressed in the ventral regions of newly formed somites (Fig. 1A). Double fluorescent in situ hybridization revealed that *bmp4^+^* somitic cells corresponded to the ventral domain of the sclerotome, marked by *nkx3.1* expression (Fig. 1B). In addition, *bmp4* expression was observed in the dorsal spinal cord and in bilateral cells along the yolk extension, adjacent to the dorsal and ventral sclerotome domains, respectively (Fig. 1B). The sclerotome gives rise to various fibroblast populations, including fin mesenchymal cells in the fin fold (Ma et al., 2023). Interestingly, at 48 hpf, *bmp2b*, *bmp4*, and *bmp6* were expressed specifically in *fras1^+^* apical epidermal cells of the fin fold and adjacent to *fbln1^+^* fin mesenchymal cells (Figs. 1C, S1A, and S1B). Together, the proximity of BMP ligand expression to the sclerotome and its derivatives suggests a crucial role for BMP signaling in sclerotome development.

**Figure 1.**
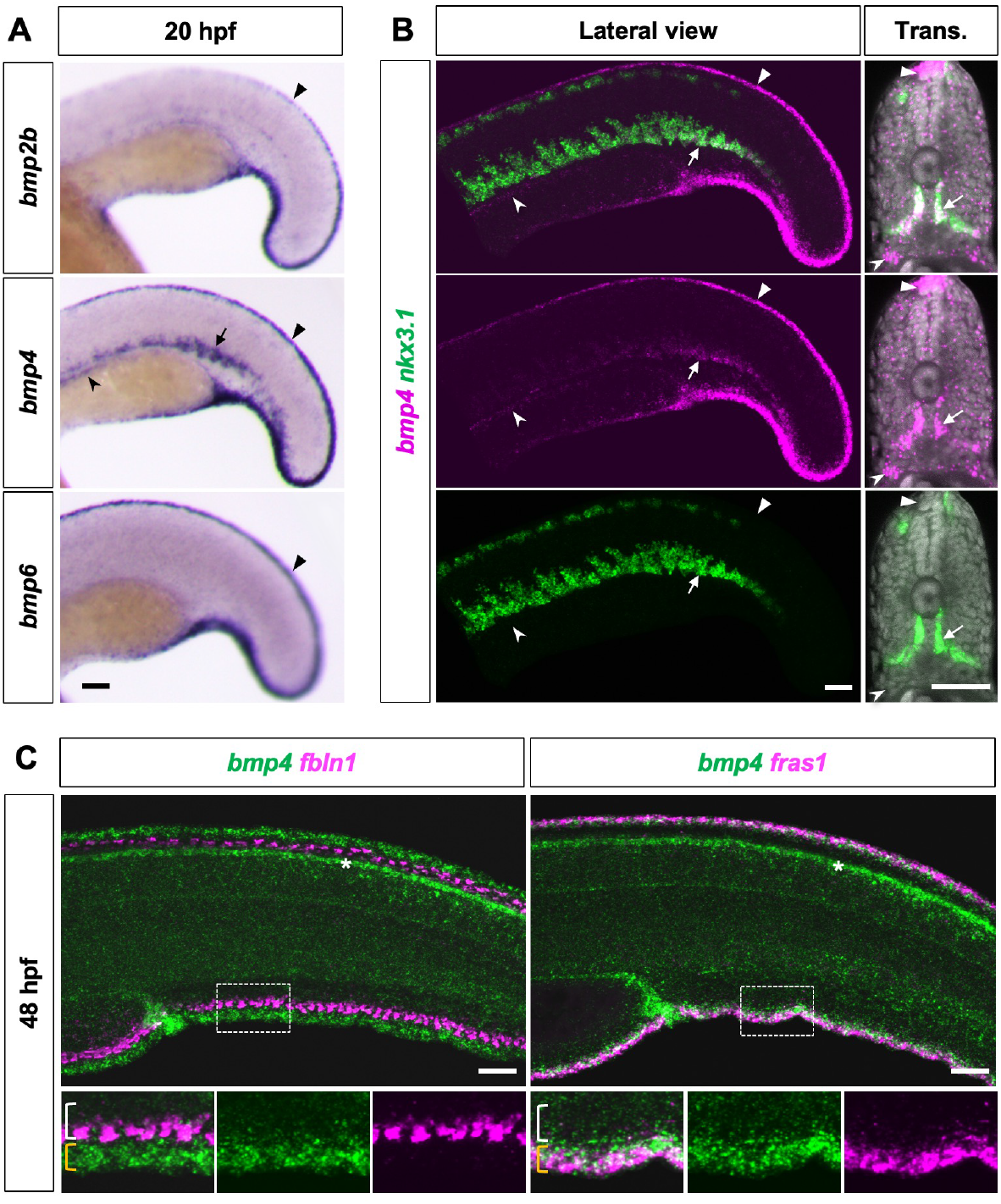
Expression analysis of BMP ligands. (A) In situ hybridization staining of BMP genes in wild-type embryos at 20 hpf in lateral views. *bmp2b*, *bmp4*, and *bmp6* are expressed along the edge of the fin fold (arrowheads). *bmp4* is also expressed in the ventral somite (arrow) as well as cells along the yolk extension (notched arrowhead). *n* = 10-15 embryos per probe. (B) Double fluorescent in situ hybridization staining of *nkx3.1* (green) and *bmp4* (magenta) in wild-type fish at 24 hpf in lateral and transverse views. *bmp4* is expressed in the *nkx3.1^+^* ventral sclerotome domain (arrows), cells ventrolateral to the ventral sclerotome (notched arrowheads), as well as the dorsal spinal cord (arrowheads). Nuclear staining using Draq5 is shown in the transverse sections. *n* = 15 embryos per staining. (C) Double fluorescent in situ hybridization staining of *bmp4* (green) with *fbln1* or *fras1* (magenta) at 48 hpf. *bmp4* is expressed in *fras1^+^* apical epidermal cells (yellow brackets) adjacent to *fbln1^+^* fin mesenchymal cells (white brackets). Asterisks denote *bmp4* expression in the dorsal spinal cord. Magnified views of boxed regions are shown in merged and individual channels. *n* = 15 embryos per staining. Scale bars: 50 μm.

### BMP signaling is necessary and sufficient for sclerotome induction

To test the role of BMP signaling in sclerotome development, we inhibited BMP signaling using *hsp70:noggin3*, whereby the expression of *noggin3*, encoding an antagonist of BMP signaling, is under the control of the heat shock inducible promoter (Chocron et al., 2007). *hsp70:noggin3* embryos or wild-type siblings were heat shocked at 18 hpf and stained with sclerotome markers *nkx3.1* and *pax9* at 24 hpf (Fig. 2A). The expression of both *nkx3.1* and *pax9* was completely lost in the somites formed after the heat shock (posterior to the yolk extension) (Fig. 2A), suggesting that BMP signaling is necessary for sclerotome induction. Interestingly, somites formed prior to the heat shock (anterior to somite 18) exhibited a loss of dorsal sclerotome domains and reduced ventral domains (Fig. 2A), suggesting that BMP signaling is also required for the maintenance of sclerotome domains. Similarly, early induction of *hsp70:noggin3* at 10 hpf resulted in the complete loss of both dorsal and ventral sclerotome domains in all somites (Fig. S2A), indicating a crucial role of BMP signaling in sclerotome induction.

**Figure 2.**
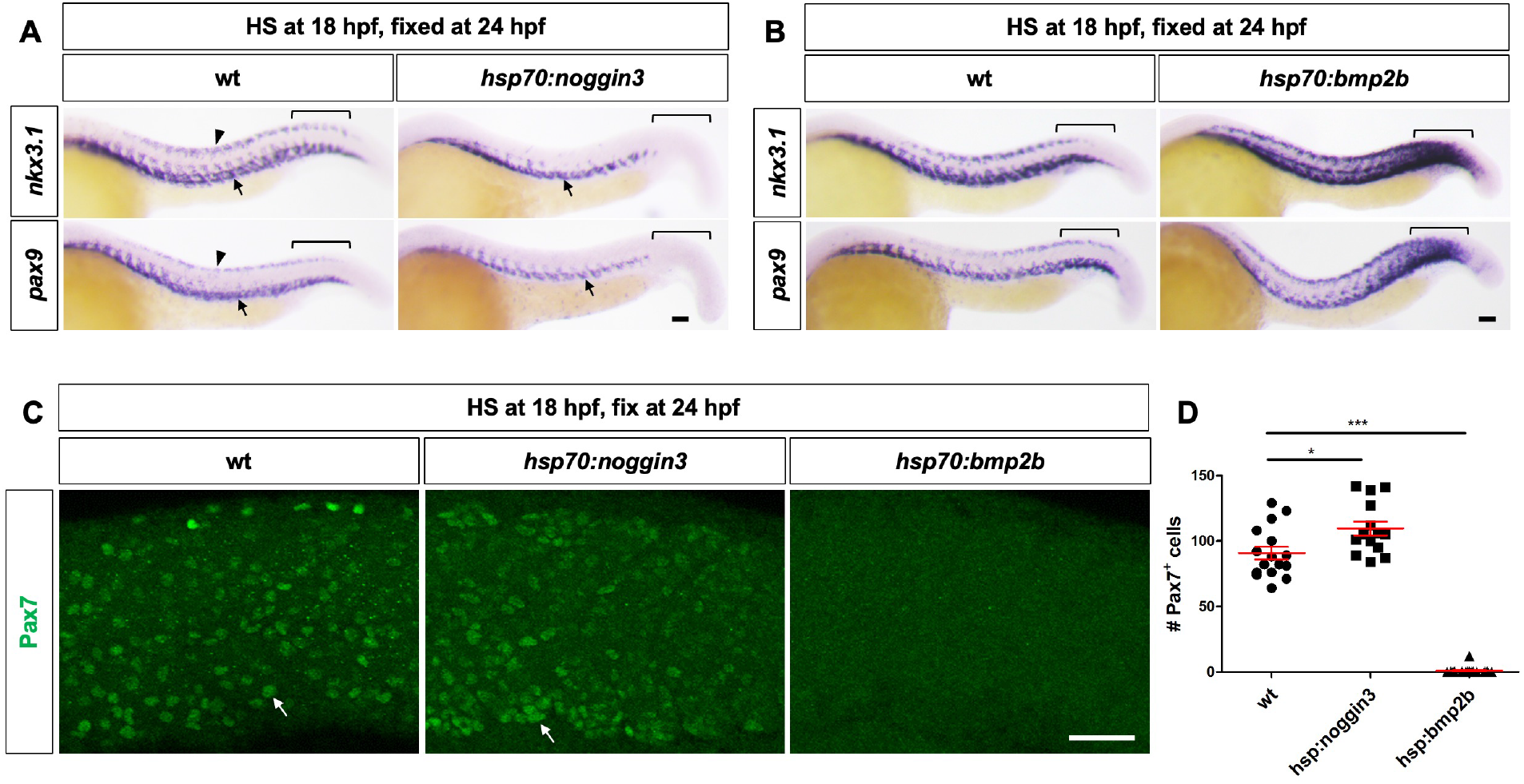
BMP signaling is necessary and sufficient for sclerotome induction. (A) Wild-type and *hsp70:noggin3* embryos were heat shocked at 18 hpf and stained at 24 hpf using *nkx3.1* and *pax9* probes. In *hsp70:noggin3* fish, the expression of *nkx3.1* and *pax9* is completely absent in the somites formed after the heat shock (brackets). In somites formed prior to the heat shock, *nkx3.1* and *pax9* expression is lost in dorsal sclerotome domains (arrowheads) and reduced in ventral sclerotome domains (arrows). *n* = 35-40 embryos per condition. (B) Wild-type and *hsp70:bmp2b* embryos were heat shocked at 18 hpf and stained at 24 hpf using *nkx3.1* and *pax9* probes. The expression of *nkx3.1* and *pax9* is expanded in *hsp70:bmp2b* fish, especially in the somites formed after the heat shock (brackets). *n* = 35-40 embryos per condition. (C) Wild-type, *hsp70:noggin3*, and *hsp70:bmp2b* embryos were heat shocked at 18 hpf and stained at 24 hpf with the Pax7 antibody. Pax7^+^ dermomyotome cells (green) are indicated by arrows. *n* = 8-15 embryos per condition. (D) Quantification of the number of Pax7^+^ cells in fish shown in (C). Each point represents the total number of Pax7^+^ dermomyotome cells in a 4-somite region of one fish. Data are plotted as mean ± SEM. Statistics: Mann-Whitney *U* test; *p* < 0.05 (*); *p* < 0.001 (***). Scale bars: 50 μm.

In converse experiments, we utilized the heat shock inducible BMP gain-of-function line, *hsp70:bmp2b*, to activate BMP signaling (Chocron et al., 2007). Compared with sibling controls, heat shock induction of *hsp70:bmp2b* at 18 hpf resulted in significantly expanded expression of *nkx3.1* and *pax9* at 24 hpf, especially in the somites formed after the heat shock (posterior to the yolk extension) (Fig. 2B). Consistent with this result, early expression of *hsp70:bmp2b* before somite formation at 10 hpf led to substantial expansion of both *nkx3.1* and *pax9* in all somites in the trunk (Fig. S2B). These results suggest that BMP signaling is also sufficient to drive sclerotome induction.

Since manipulation of BMP signaling alters the size of the sclerotome compartment in the somite, we predict opposite changes in other somitic compartments. Specifically, we examined the dermomyotome, which can be identified by Pax7 antibody staining. Inhibition of BMP signaling by *hsp70:noggin3* resulted in significantly more Pax7^+^ dermomyotome cells, while BMP activation using *hsp70:bmp2b* led to a complete loss of Pax7 staining (Fig. 2C and 2D). Therefore, the level of BMP signaling likely controls the relative proportion of different compartments within the somite.

### Inhibition of BMP signaling results in the loss of sclerotome-derived cells

Our results suggest that BMP signaling is required for sclerotome induction. We predict that early inhibition of BMP signaling will result in the loss of cells derived from the sclerotome lineage, including tenocytes and fin mesenchymal cells (Ma et al., 2023). To test this, we induced *hsp70:noggin3* expression at 18 hpf and stained fish with two tenocyte markers, *tnmd* and *prelp*, at 54 hpf. Compared to their sibling controls, expression of both tenocyte markers in the tail region (formed after the heat shock) was completely lost, and tenocytes along the dorsal myotendinous junction in anterior regions were also absent (Fig. 3A). This pattern of tenocyte loss is consistent with the observed reduction in sclerotome domains following BMP signaling inhibition (Fig. 2A). Similar to tenocytes, inhibition of BMP signaling by DMH1, a BMP receptor inhibitor (Hao et al., 2010), from 18 to 45.5 hpf, resulted in reduced *fbln1^+^* fin mesenchymal cells in both dorsal and ventral fin folds (Fig. 3B). To quantify the number of fin mesenchymal cells, we treated *kdrl:EGFP*; *nkx3.1:Gal4; UAS:NTR-mCherry* (the Gal4/UAS transgenic lines are designated as *nkx3.1^NTR-mCherry^*) embryos with DMH1 from 18 to 48 hpf. The DMH1-treated fish showed poorly formed caudal vein plexus (CVP) (Fig. 3C), a structure dependent on active BMP signaling (Wiley and Jin, 2011), demonstrating the efficacy of DMH1 as a BMP signaling inhibitor. Strikingly, DMH1 treatment led to a significant reduction in the number of mCherry^+^ fin mesenchymal cells in both the dorsal (49% reduction) and ventral (63% reduction) fin folds (Fig. 3C and 3D). Together, our results suggest that inhibition of BMP signaling impairs sclerotome induction, leading to the loss of sclerotome-derived cell lineages.

**Figure 3.**
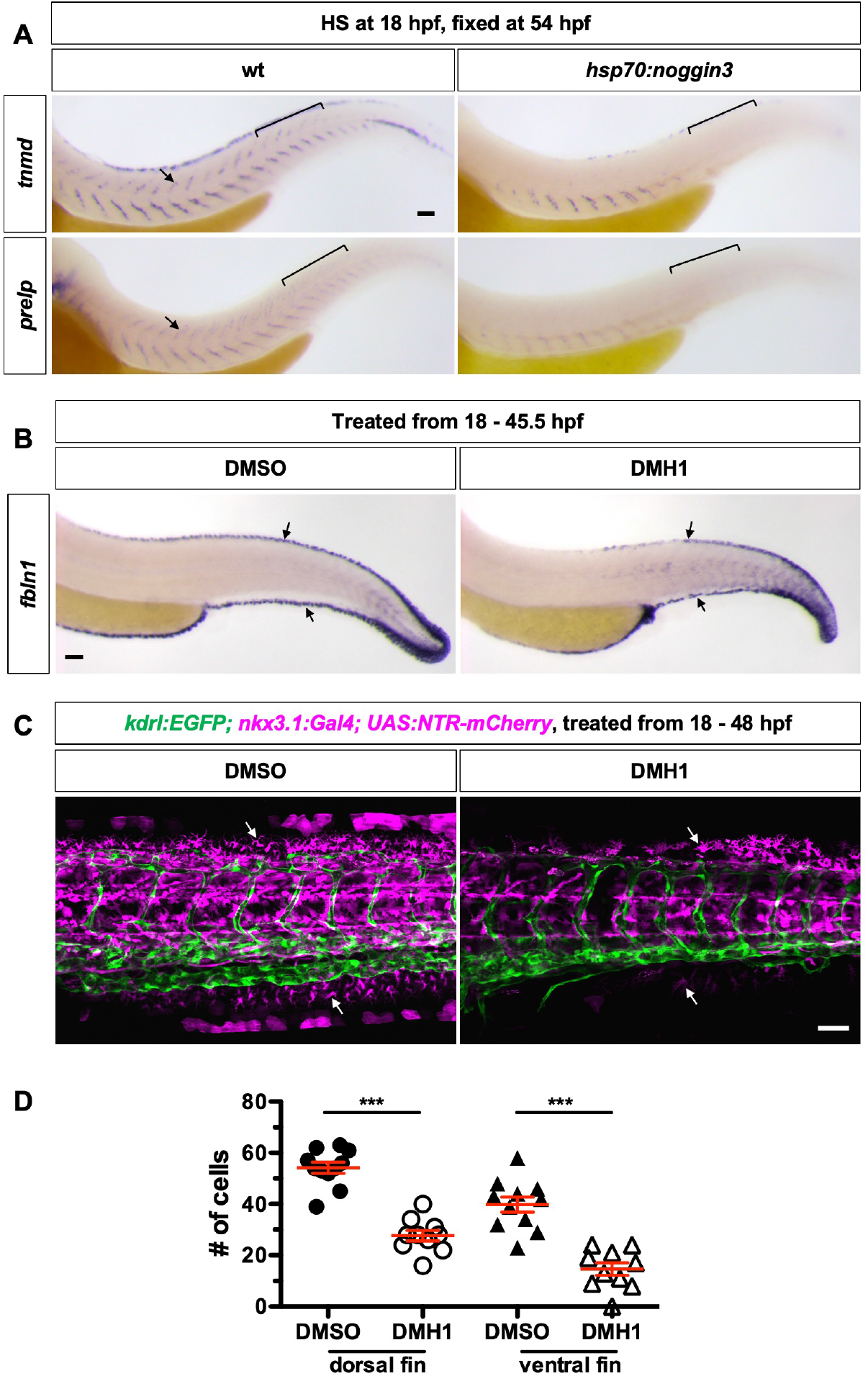
Inhibition of BMP signaling results in the loss of sclerotome-derived cells. (A) Wild-type and *hsp70:noggin3* fish were heat shocked at 18 hpf and stained at 54 hpf using *tnmd* and *prelp* probes. Compared to wild-type controls, the number of *tnmd^+^* and *prelp^+^* tenocytes (arrows) along the MTJ is greatly reduced in *hsp70:noggin3* embryos, especially in somites formed after the heat shock (brackets). *n* = 8-15 embryos per condition. (B) Wild-type embryos were treated with DMSO or DMH1 from 18 to 45.5 hpf and stained with the *fbln1* probe. *fbln1* expression is markedly reduced in both the dorsal and ventral fin folds (arrows) compared to DMSO controls. *n* = 15 embryos per condition. (C) *kdrl:EGFP; nkx3.1:Gal4; UAS:NTR-mCherry* embryos were treated with either DMSO or DMH1 between 18 and 48 hpf. DMH1 treatment resulted in a reduction of mCherry^+^ fin mesenchymal cells (arrows) in both the dorsal and ventral fin folds compared to DMSO-treated controls. *n* = 19 embryos per condition. (D) Quantification of the number of fin mesenchymal cells. Fin mesenchymal cells were counted in either the dorsal or ventral fin fold between somites 18 and 28. Each point represents the total number of fin mesenchymal cells in a 10-somite region of one fish. *n* = 11 (DMSO) and 10 (DMH1) embryos. Data are plotted as mean ± SEM. Statistics: Mann-Whitney *U* test; *p* < 0.001 (***). Scale bars: 50 μm.

### BMP signaling functions cell-autonomously in sclerotome induction

Given that BMP signaling is required for sclerotome induction, we asked if BMP signaling is active in sclerotome progenitors. We used phospho-Smad1/5/9 (pSmad) staining as a sensitive readout for BMP signaling activity. In *nkx3.1^NTR-mCherry^* embryos at 24 hpf, pSmad staining was detected in various levels in many somitic cells, including mCherry^+^ sclerotome progenitors in both the dorsal and ventral domains (Fig. 4A). In contrast, induction of *hsp70:noggin3* completely abolished both pSmad and *nkx3.1^NTR-mCherry^* expression in the somite (Fig. 4A). By normalizing to the pSmad level in *hsp70:noggin3* embryos, we quantified pSmad intensity in mCherry^+^ sclerotome progenitors in wild-type embryos. On average, sclerotome cells in both the dorsal and ventral domains displayed significantly higher pSmad intensity compared to *hsp70:noggin3* controls (Fig. 4B), suggesting active BMP signaling in sclerotome progenitors.

**Figure 4.**
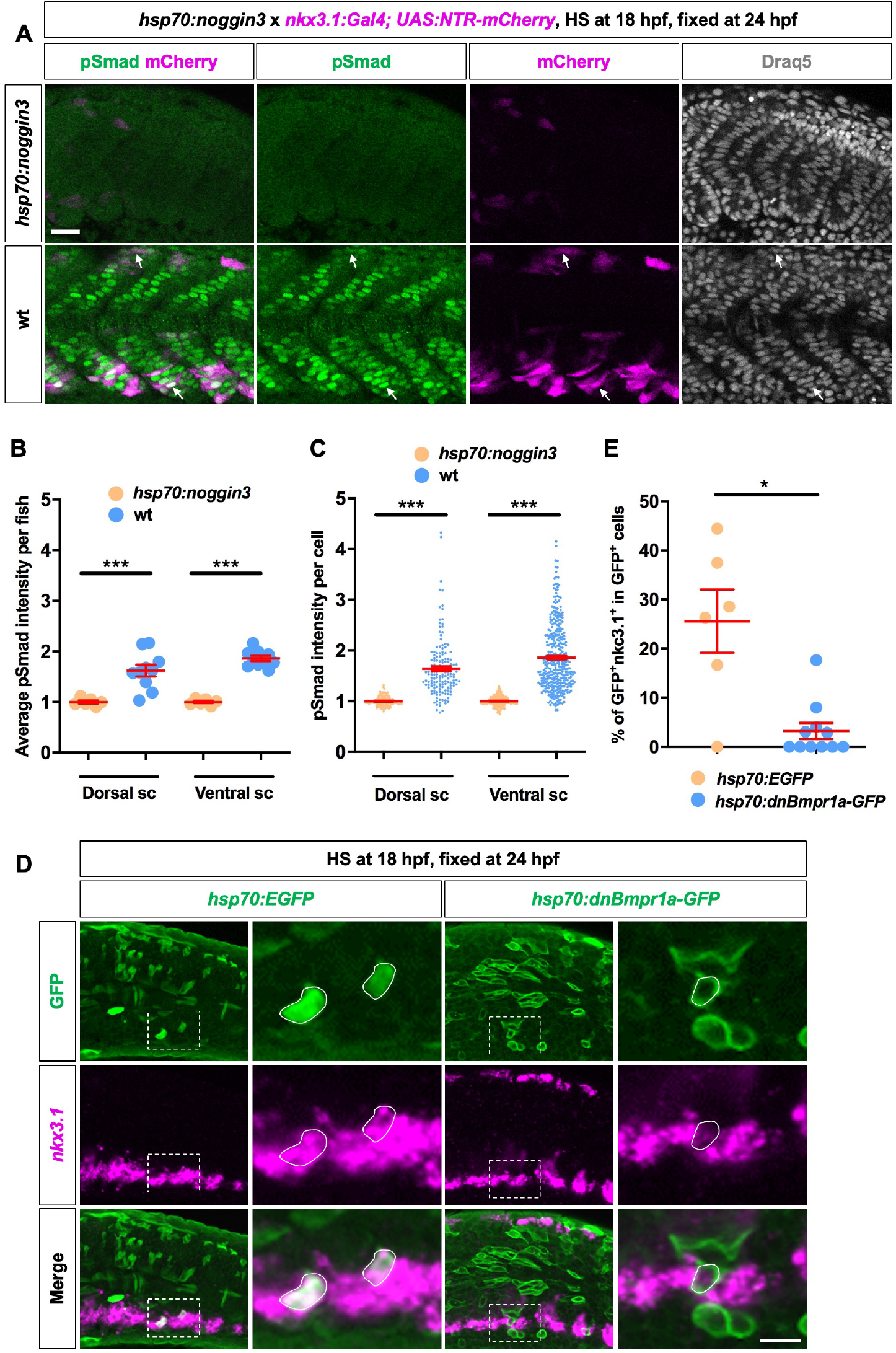
BMP signaling regulates sclerotome induction in a cell-autonomous manner. (A) Embryos from a cross between *hsp70:noggin3* and *nkx3.1:Gal4; UAS:NTR-mCherry* were heat shocked at 18 hpf and stained at 24 hpf with the pSmad antibody (green) and the nuclear dye Draq5 (grey). In wild-type fish, pSmad^+^mCherry^+^ cells (arrows) are present in both dorsal and ventral sclerotome domains, whereas pSmad and mCherry expression are completely lost in *hsp70:noggin3* embryos. *n* = 15 embryos per condition. (B-C) Quantification of the pSmad intensity in dorsal and ventral sclerotome cells in fish shown in (A). pSmad intensity was measured in mCherry^+^ cells of wild-type fish or cells in dorsal/ventral somites in *hsp70:noggin3* embryos in somites 18-22 (formed after the heat shock). Each point represents the average pSmad intensity of one fish (B) or pSmad intensity of one cell (C), both of which are normalized to that of *hsp70:noggin3* embryos. *n* = 6 (*hsp70:noggin3*) and 10 (wild-type) fish. (D) Wild-type embryos were injected with *hsp70:EGFP* or *hsp70:dnBmpr1a-GFP* plasmids, heat shocked at 18 hpf, and stained at 24 hpf using the GFP antibody (green) and *nkx3.1* probe (magenta). EGFP^+^ cells in the ventral somite express *nkx3.1* normally, whereas dnBmpr1a-GFP^+^ cells show minimal *nkx3.1* expression despite being located in the *nkx3.1^+^* ventral sclerotome domain. Magnified views of boxed regions are shown with labeled cells of interest outlined in white. *n* = 6 (*hsp70:EGFP*) and 11 (*hsp70:dnBmpr1a-GFP*) fish. (E) Quantification of the percentage of *nkx3.1*-expressing cells out of all GFP^+^ cells in the dorsal and ventral somites in the experiment shown in (D). Data in (B, C, and E) are plotted as mean ± SEM. Statistics: Mann-Whitney *U* test; *p* < 0.05 (*); *p* < 0.01 (**); *p* < 0.001 (***). Scale bars: 50 μm.

Interestingly, the distribution of pSmad intensity among individual sclerotome cells exhibited considerable heterogeneity (Fig. 4C), which might reflect the transient nature of BMP response. Our results suggest that most sclerotome progenitors in wild-type embryos show active BMP signaling.

Next, we asked if BMP signaling is required cell-autonomously in inducing the sclerotome fate. We utilized a heat shock inducible construct expressing dominant-negative BMP receptor, *hsp70:dnBmpr1a-GFP*, in which the intracellular kinase domain of *Xenopus* BMP receptor 1a is replaced with GFP (Pyati et al., 2005). As a control, we used *hsp70:EGFP*, which expresses EGFP under the control of the heat shock promoter. Wild-type embryos were injected with either *hsp70:EGFP* or *hsp70:dnBmpr1a-GFP*, heat shocked at 18 hpf, and co-stained with the *nkx3.1* probe and the GFP antibody. In *hsp70:EGFP*-injected controls, 25.6% of GFP^+^ cells in the dorsal or ventral somite expressed sclerotome marker *nkx3.1* at a level comparable to their neighboring GFP^-^ cells, suggesting normal induction of the sclerotome fate (Fig. 4D and 4E). By contrast, in *hsp70:dnBmpr1a-GFP*-injected embryos, most dnBmpr1a-GFP^+^ cells lacked any *nkx3.1* expression even though their GFP^-^ neighbors in the sclerotome domain showed robust *nkx3.1* expression (Fig. 4D and 4E). Together, these results suggest that BMP signaling induces the sclerotome fate in a cell-autonomous manner.

### Sclerotome-derived cells show distinct BMP response

Since BMP signaling is required for inducing sclerotome progenitors, we asked if BMP signaling persists in cells derived from the sclerotome. Using *nkx3.1^NTR-mCherry^* to label sclerotome-derived cells, we performed pSmad staining in embryos at 2 dpf. Strikingly, mCherry^+^ cells around the notochord showed no pSmad staining, while mCherry^+^ fin mesenchymal cells in the fin fold exhibited robust pSmad expression (Fig. 5A). To confirm this finding further, we crossed *nkx3.1^NTR-mCherry^* with *BRE:GFP*, a live reporter of BMP response (Collery and Link, 2011). At 2 dpf, mCherry^+^ notochord-associated cells were largely GFP^-^, suggesting the lack of active BMP signaling (Fig. 5B). By contrast, mCherry^+^ fin mesenchymal cells displayed strong *BRE:GFP* expression, indicative of active BMP response (Fig. 5B). These results suggest that different sclerotome-derived cells display distinct BMP response: cells in the medial trunk turn off BMP response, whereas fin mesenchymal cells in the fin fold maintain active BMP signaling. Interestingly, BMP-responding fin mesenchymal cells are located adjacent to BMP ligand-expressing apical epidermal cells (Figs. 1C and S1), suggesting paracrine signaling between these two cell layers.

**Figure 5.**
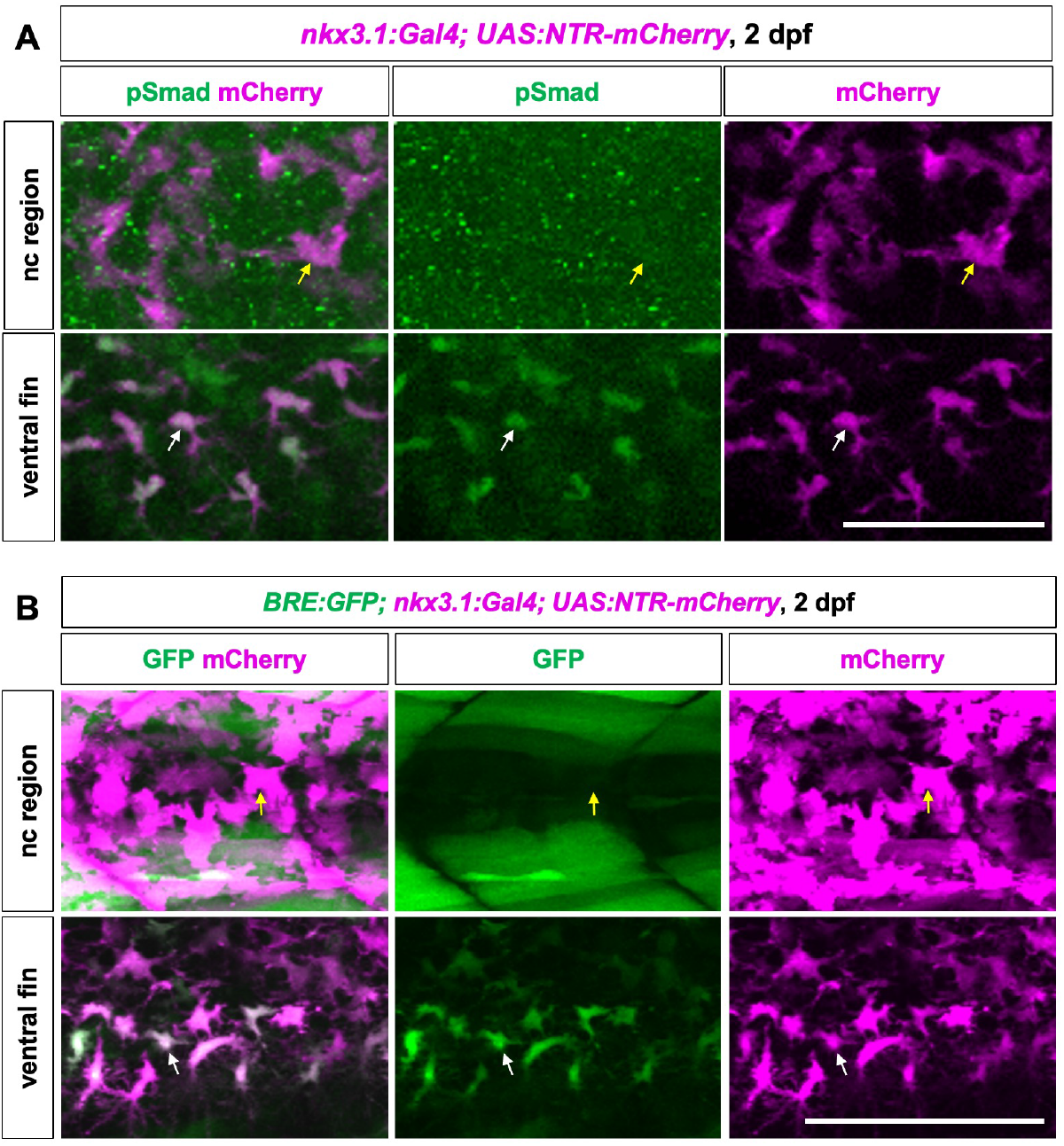
Sclerotome-derived cells show distinct BMP response. (A) *nkx3.1:Gal4; UAS:NTR-mCherry* embryos were stained with the pSmad antibody (green) at 2 dpf. mCherry^+^ sclerotome-derived cells (yellow arrows) around the notochord (nc) show no pSmad staining, whereas mCherry^+^ fin mesenchymal cells (while arrows) are pSmad^+^. *n* = 14 embryos. (B) *nkx3.1:Gal4; UAS:NTR-mCherry* fish were crossed to *BRE:GFP* to label BMP-responding cells at 2 dpf. mCherry^+^ fin mesenchymal cells (white arrows) in the ventral fin fold are marked by strong GFP expression. In contrast, mCherry^+^ sclerotome-derived cells (yellow arrows) around the notochord are GFP^-^. *n* = 9 embryos. Scale bars: 50 μm.

### Downregulation of BMP signaling is required for the differentiation of sclerotome-derived cells in the medial trunk

The lack of BMP response in sclerotome-derived notochord-associated cells suggests that active BMP signaling is not required in these cells. Consistent with this prediction, inhibition of BMP signaling by *hsp70:noggin3* at 18 hpf did not alter the expression of *pax1a* and *nkx3.2*, two markers of sclerotome-derived cells around the notochord, in the anterior somites (formed before 18 hpf) at 24 hpf (Fig. 6A). Surprisingly, activation of BMP signaling by *hsp70:bmp2b* at 18 hpf completely abolished the expression of both *pax1a* and *nkx3.2* (Fig. 6B), suggesting that sustained active BMP signaling in sclerotome-derived cells interferes with their proper differentiation. In line with this finding, induction of *hsp70:bmp2b* expression at 24 hpf, a stage characterized by *pax1a* and *nkx3.2* expression in sclerotome-derived cells around the notochord (Ma et al., 2018), also resulted in the complete loss of both marker expression at 30 hpf (Fig. 6C). Similarly, sustained BMP signaling activation led to the complete loss of expression of *ola-twist1:EGFP* (Fig. 6D), a transgenic reporter for sclerotome-derived notochord-associated cells (Ma et al., 2018).

**Figure 6.**
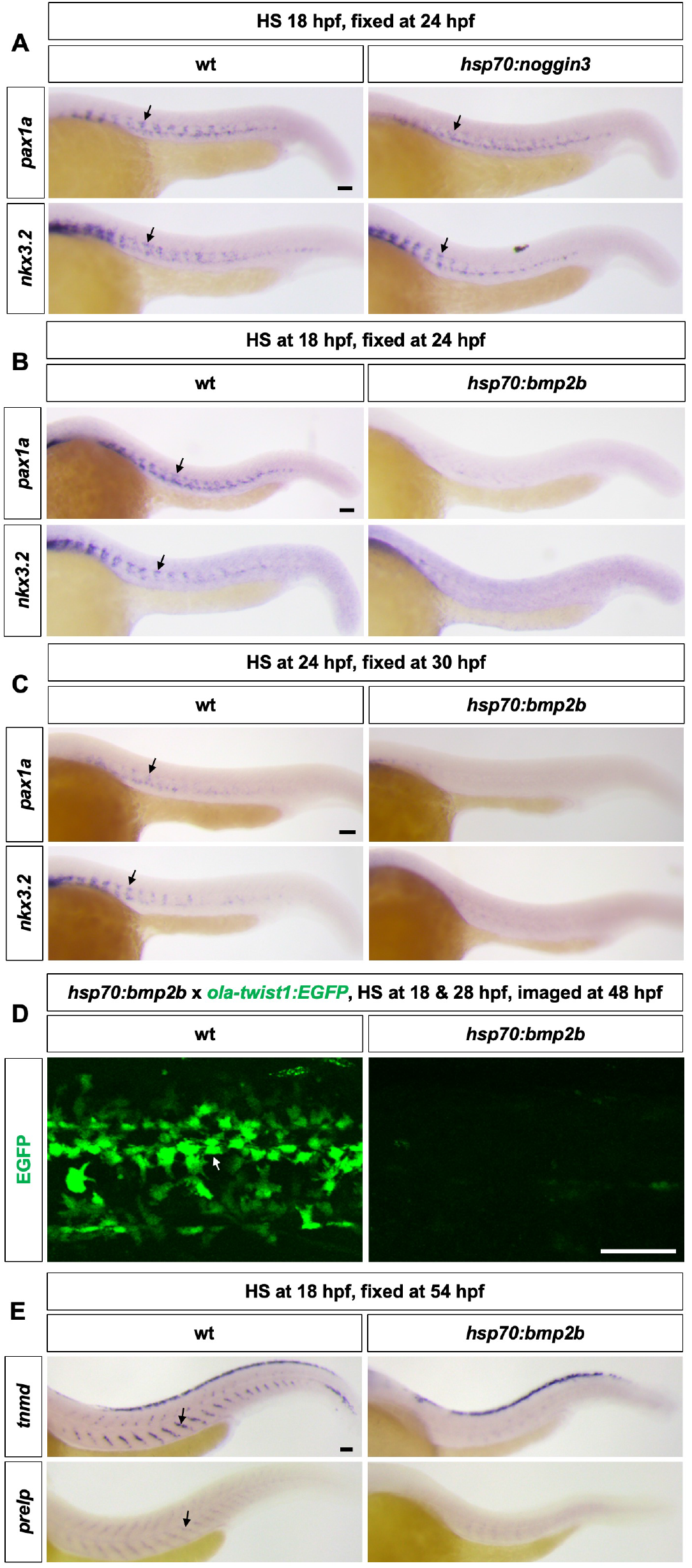
Downregulation of BMP signaling is required for the differentiation of sclerotome-derived cells. (A) Wild-type and *hsp70:noggin3* embryos were heat shocked at 18 hpf and stained with *pax1a* and *nkx3.2* probes at 24 hpf. The expression of *pax1a* and *nkx3.2* around the notochord in the anterior regions (arrows) is largely normal in *hsp70:noggin3* fish. *n* = 12 embryos per condition. (B) Wild-type and *hsp70:bmp2b* embryos were heat shocked at 18 hpf and stained with *pax1a* and *nkx3.2* probes at 24 hpf. Compared to controls, *pax1a* and *nkx3.2* expression around the notochord (arrows) is completely absent in *hsp70:bmp2b* embryos. *n* = 12 embryos per condition. (C) Wild-type and *hsp70:bmp2b* embryos were heat shocked at 24 hpf and stained with *pax1a* and *nkx3.2* probes at 30 hpf. The expression of *pax1a* and *nkx3.2* is completely lost in *hsp70:bmp2b* embryos. *n* = 12 embryos per condition. (D) Embryos from a cross between *hsp70:bmp2b* and *ola-twist1:EGFP* fish were heat shocked at 18 and 28 hpf, and imaged at 48 hpf. Expression of *ola-twist1:EGFP* in notochord-associated sclerotome-derived cells (arrow) is completely lost in *hsp70:bmp2b* embryos compared to the wild-type controls. *n* = 8 embryos per group. (E) Wild-type and *hsp70:bmp2b* embryos were heat shocked at 18 hpf and stained with *tnmd* and *prelp* probes at 54 hpf. Compared to controls, *tnmd^+^*and *prelp^+^* tenocytes (arrows) are largely absent in *hsp70:bmp2b* fish. *n* = 12 embryos per condition. Scale bars: 50 μm.

Since the sclerotome gives rise to different types of fibroblasts, including tenocytes (Ma et al., 2018; Ma et al., 2023), we examined the impact of sustained BMP activation on tenocyte differentiation. Similar to sclerotome-derived cells around the notochord, prolonged activation of BMP signaling resulted in a loss of expression of both *tnmd* and *prelp* (Fig. 6E). Together, our results suggest that although BMP signaling is required for sclerotome induction, BMP response needs to be downregulated to allow the proper differentiation of sclerotome-derived cells in the medial trunk.

### Active BMP signaling is required for the proper morphogenesis of fin mesenchymal cells

In contrast to sclerotome-derived notochord-associated cells, fin mesenchymal cells show active BMP response (Fig. 5). To test if BMP signaling is required for the development of fin mesenchymal cells, we blocked BMP signaling after the sclerotome is induced in the posterior trunk. We treated *kdrl:EGFP; nkx3.1^NTR-mCherry^* embryos with DMSO or DMH1 from 25 to 49 hpf. Compared to the DMSO-treated controls, DMH1-treated embryos exhibited significantly expanded fin folds (Fig. 7A and 7B). Nonetheless, the number of fin mesenchymal cells remained largely unaffected in DMH1-treated fish (Fig. 7C). Strikingly, DMH1-treated fin mesenchymal cells displayed a hyperbranching morphology with significantly more cellular projections compared to DMSO-treated controls (Fig. 7A and 7D). This result suggests that active BMP signaling is required for the proper morphogenesis of fin mesenchymal cells during development.

**Figure 7.**
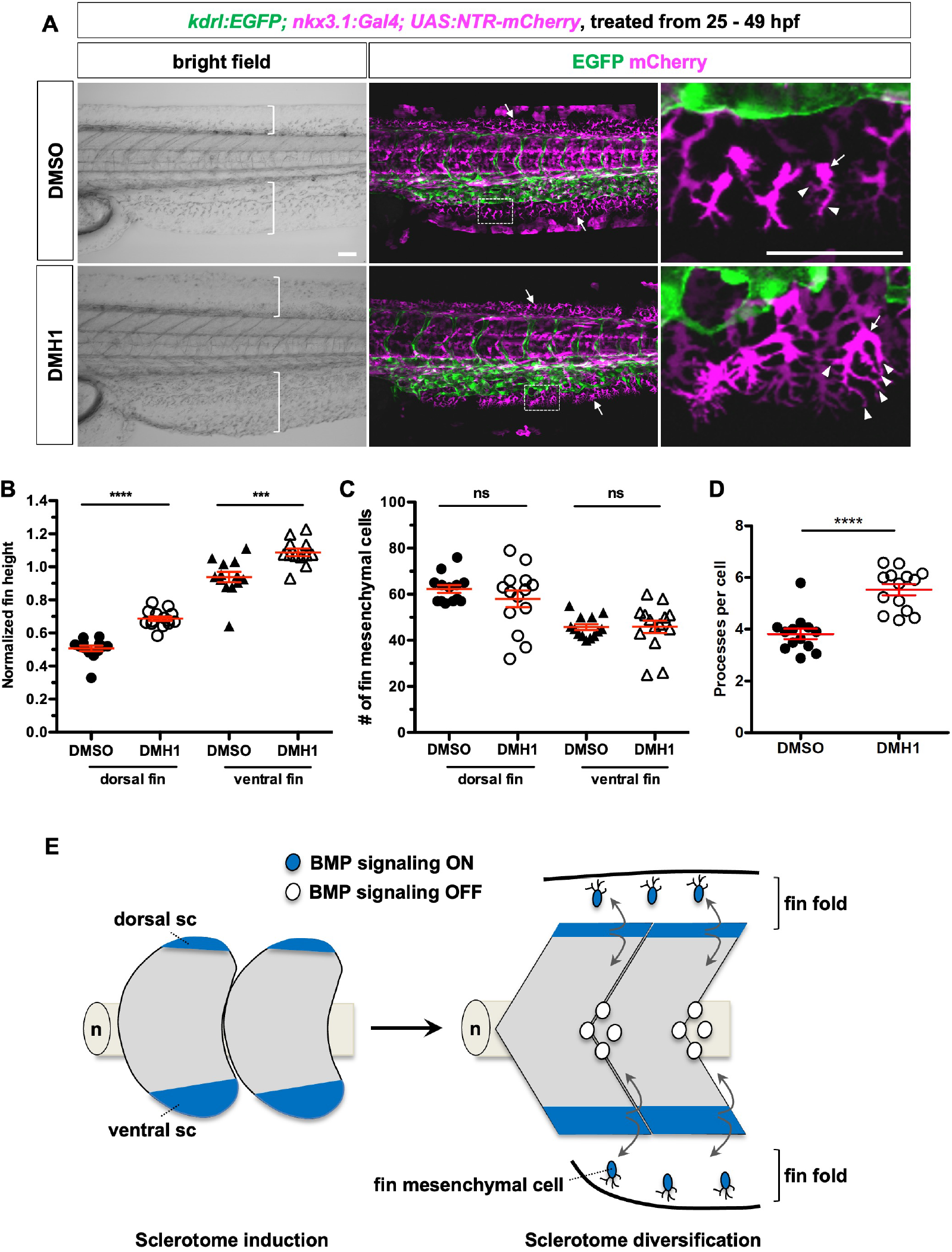
BMP signaling regulates the morphogenesis of fin mesenchymal cells. (A) *kdrl:EGFP; nkx3.1:Gal4; UAS:NTR-mCherry* embryos were treated with either DMSO or DMH1 from 25 to 49 hpf. DMH1 treatment results in the expansion of both dorsal and ventral fin folds (brackets). Fin mesenchymal cells (arrows) in DMH1-treated embryos have more branches (arrowheads) compared to DMSO-treated controls. Magnified views of boxed regions are shown. *n* = 13 (DMSO) and 14 (DMH1) embryos. (B) Quantification of the fin fold expansion in DMSO/DMH1-treated embryos. The heights of the dorsal fin fold, ventral fin fold, and somite were measured at somite 23. Fin fold height was normalized to the height of the somite of the same embryo. (C) Quantification of the number of fin mesenchymal cells in DMSO/DMH1-treated fish. Fin mesenchymal cells were counted in either the dorsal or ventral fin fold between somites 18 and 28. Each point represents the total number of fin mesenchymal cells in one fish. (D) Quantification of the number of processes of fin mesenchymal cells. The processes of fin mesenchymal cells were counted in the ventral fin fold between somites 18 and 28. Each point represents the average number of processes of one fin mesenchymal cell. (E) Model of dynamic BMP signaling in regulating zebrafish sclerotome development. In newly formed somites, active BMP signaling is required for the induction of sclerotome (sc) domains at the dorsal and ventral regions of the somite. During sclerotome lineage diversification, attenuation of BMP signaling in sclerotome-derived cells surrounding the notochord (n) is essential for their differentiation, whereas sustained active BMP signaling is crucial for the morphogenesis of fin mesenchymal cells in the fin fold. Active BMP signaling is indicated in blue for cells of the sclerotome lineage. All data are plotted as mean ± SEM. Statistics: Mann-Whitney *U* test; *p* > 0.05 (ns, not significant); *p* < 0.001 (***); *p* < 0.0001 (****). Scale bars: 50 μm.

## DISCUSSION

Our work reveals a dynamic role of BMP signaling in the development of the sclerotome lineage in zebrafish (Fig. 7E). First, active BMP signaling is required for sclerotome induction in a cell-autonomous manner. Second, downregulation of BMP signaling is essential for the development of sclerotome-derived notochord-associated cells as well as tenocytes. Third, sustained BMP signaling is required for the proper morphogenesis of fin mesenchymal cells.

### Active BMP signaling is required for sclerotome induction

Studies in chick and mouse suggest that active Hedgehog (Hh) signaling is critical for early sclerotome development (Christ et al., 2004; Monsoro-Burq, 2005). Interestingly, our previous work in zebrafish shows that sclerotome domains can form normally even in the absence of active Hh signaling (Ma et al., 2018), suggesting that alternative signals are crucial for sclerotome induction. Our current work provides three lines of evidence indicating that active BMP signaling is essential for inducing the sclerotome compartment. First, BMP ligands are expressed in or near the sclerotome domains: *bmp4* is transiently expressed in the ventral sclerotome domain, while the dorsal sclerotome domain is adjacent to *bmp4*-expressing dorsal spinal cord. Second, most sclerotome progenitors in both the dorsal and ventral domains exhibit active BMP response, as indicated by elevated pSmad staining. Third, gain- and loss-of-function experiments demonstrate that BMP signaling is both necessary and sufficient to drive sclerotome marker expression. Importantly, mosaic inhibition of BMP signaling reveals that BMP signaling acts cell-autonomously in inducing the sclerotome fate.

Our findings contrast with the general model in amniotes, where inhibition of BMP signaling is thought to be required for sclerotome induction (Stafford et al., 2011). In mice, broad activation of BMP signaling by genetically removing two BMP antagonists, NOGGIN and GREMLIN1, in *Noggin; Gremlin1* double mutants results in a complete loss of expression of sclerotome markers, including *Pax1* (Stafford et al., 2011). However, grafting experiments in chick suggest that different regions of the sclerotome exhibit distinct requirement for BMP signaling (Christ et al., 2004). The central and ventral sclerotome, marked by *Pax1* expression, is controlled by Shh and Noggin, whereas the dorsal and lateral regions of the sclerotome, which lack *Pax1* expression, depend on BMP-4 (Ebensperger et al., 1995; Monsoro-Burq et al., 1996; Monsoro-Burq and Le Douarin, 2000; Sudo et al., 2001). Grafting BMP-producing cells near the dorsal sclerotome promotes the formation of dorsal mesenchyme and enlarged dorsal vertebral cartilages (Monsoro-Burq et al., 1996; Monsoro-Burq and Le Douarin, 2000). By contrast, grafting BMP-producing cells near the ventral sclerotome inhibits *Pax1* expression, resulting in reduced vertebral bodies (Monsoro-Burq et al., 1996; Hirsinger et al., 1997; Monsoro-Burq and Le Douarin, 2000). Interestingly, in zebrafish, we find that *pax1a^+^*sclerotome-derived notochord-associated cells require active Hh signaling (Ma et al., 2018), while sustained activation of BMP signaling in these cells results in a complete loss their marker expression. Based on marker expression and BMP/Hh regulation, we suggest that the dorsal and lateral *Pax1^-^*sclerotome in chick likely corresponds to the dorsal and ventral sclerotome domains in zebrafish, respectively, whereas the central/ventral *Pax1^+^*sclerotome in chick is the equivalent of *pax1a^+^* sclerotome-derived notochord-associated cells in zebrafish. Therefore, in both zebrafish and amniotes, BMP signaling plays an active role in inducing the *Pax1^-^*sclerotome domains, whereas inhibition of BMP signaling is required for the development of the *Pax1^+^*sclerotome regions.

### Differential regulation of BMP signaling is essential for sclerotome lineage diversification

The zebrafish sclerotome gives rise to diverse fibroblast subtypes that are widely distributed throughout the trunk (Ma et al., 2023). Although sclerotome progenitors are induced by active BMP signaling, cells derived from the sclerotome lineage exhibit distinct BMP responses in a location-dependent manner. In the periphery, fin mesenchymal cells in the fin fold display prominent BMP pathway activation. By contrast, sclerotome-derived cells around the notochord show an absence of BMP signaling. The local signaling microenvironment likely determines whether sclerotome-derived cells respond to BMP signaling. BMP ligands, including *bmp2b*, *bmp4*, and *bmp6*, are expressed in the apical epidermal cells of the fin fold, which are likely responsible for activating BMP signaling in the adjacent fin mesenchymal cells. Similarly, the absence of BMP response in sclerotome-derived notochord-associated cells is likely due to the combined effect of the expression of the BMP antagonist *noggin2* in the notochord (Fürthauer et al., 1999) and the lack of BMP ligand expression in the vicinity. Our timed genetic manipulations of BMP signaling reveal that temporal regulation of BMP response is functionally important for the development of different cell types from the sclerotome lineage. Sustained activation of BMP signaling results in the loss of *pax1a* and *nkx3.2* expression, specific markers for sclerotome-derived notochord-associated cells, leading to defects in the differentiation of tenocytes. Similarly, inhibition of BMP signaling compromises the proper morphogenesis of fin mesenchymal cells. Together, our results suggest that dynamic regulation of BMP signaling is essential for the diversification of the sclerotome lineage. Similar modes of temporal regulation of BMP signaling have been observed in other developmental contexts. For instance, during vertebrate neural development, BMP signaling is initially repressed to induce the neural plate from the ectoderm (Bond et al., 2012). Subsequently, active BMP signaling is necessary for the induction of neural crest cells and the patterning of the dorsal spinal cord (Bond et al., 2012).

### BMP signaling regulates the morphogenesis of fin mesenchymal cells

Our studies reveal that BMP signaling is required at multiple steps in the generation of fin mesenchymal cells. Early inhibition of BMP signaling results in a reduced number of fin mesenchymal cells due to compromised sclerotome induction. By contrast, late inhibition of BMP response does not alter the number of fin mesenchymal cells but results in cells with an aberrant hyperbranching morphology. This result suggests that BMP signaling is not required for the migration of fin mesenchymal precursors into the fin fold but is necessary for their proper morphogenesis. There are two non-mutually exclusive explanations for cell morphology phenotype. First, BMP signaling might directly regulate the formation of cellular protrusions. Previous studies have implicated BMP signaling in modulating the cytoskeletal network (Foletta et al., 2003; Lee-Hoeflich et al., 2004; Podkowa et al., 2010) and the dynamics of cell processes (von der Hardt et al., 2007; Wakayama et al., 2015). Alternatively, the hyperbranching phenotype might be secondary to an altered extracellular environment due to BMP inhibition. Interesting, blockage of BMP signaling also results in a significant outgrowth of the fin fold, caused by increased cell proliferation, altered cell division orientation, and excessive distal cell migration (Ka et al., 2020). This result suggests that BMP signaling plays an additional role in restricting the outgrowth of the fin fold. It is plausible that the altered extracellular environment in the expanded fin fold upon BMP inhibition might change the morphology of fin mesenchymal cells, as previously suggested (Feitosa et al., 2012; Mahabaleshwar et al., 2022).

In summary, our work demonstrates multiple roles of BMP signaling in the diversification of the sclerotome lineage in zebrafish (Fig. 7E). Early BMP signaling is required for the induction of the sclerotome compartment in the somite. Downregulation of BMP signaling is essential for the development of sclerotome-derived cells in the medial trunk, such as cells around the notochord and tenocytes, whereas sustained activation of BMP signaling is required for the morphogenesis of fin mesenchymal cells in the fin fold. Insights from our work provide a framework to help optimize in vitro differentiation procedures for producing tissue support cells in regenerative medicine.

## MATERIALS AND METHODS

### Ethics statement

All animal research was conducted in accordance with the principles outlined in the current Guideline of the Canadian Council on Animal Care. All protocols were approved by the Animal Care Committee at the University of Calgary (#AC21-0102).

### Zebrafish strains

Zebrafish strains used in this research were raised under standard conditions. The following transgenic strains were used in our study: *Tg(BRE:GFP)mw29* (Collery and Link, 2011), *Tg(hsp70:bmp2b)fr13* (Chocron et al., 2007), *Tg(hsp70:noggin3)fr14* (Chocron et al., 2007), *Tg(kdrl:EGFP)la163* (Choi et al., 2007), *TgBAC(nkx3.1:Gal4)ca101* (Ma et al., 2018), *Tg(ola-twist1:EGFP)ca104* (Ma et al., 2018), and *Tg(UAS:NTR-mCherry)c264* (Davison et al., 2007).

### Plasmid injection

The *hsp70:dnBmpr1a-GFP* construct was generated in a Tol2 vector based on a previously published *hsp70l:dnXla.Bmpr1a-GFP* plasmid (Pyati et al., 2005). 40 pg of the *hsp70:EGFP* or *hsp70:dnBmpr1a-GFP* plasmid was co-injected with 40 pg of *tol2* transposase mRNA into wild-type embryos at the one-cell stage.

### In situ hybridization and immunohistochemistry

Whole-mount in situ hybridization and immunohistochemistry were performed according to standard protocols. We used the following RNA probes in this study: *bmp2b*, *bmp4*, *bmp6*, *fbln1*, *fras1*, *nkx3.1*, *nkx3.2, pax1a*, *pax9*, *prelp*, and *tnmd*. Double fluorescent in situ hybridizations were performed using various combinations of digoxigenin (DIG) or dinitrophenyl (DNP) labeled probes. For immunohistochemistry, the following primary antibodies were used: mouse monoclonal antibody to Pax7 (1:5, Developmental Studies Hybridoma Bank (DSHB), PAX7-supernatant), rabbit polyclonal antibody to phosphoSmad 1/5/9 (1:100, Cell Signaling Technology, 13820), and chick polyclonal antibody to GFP (1:250, Aves, GFP-1020). The appropriate Alexa Fluor-conjugated secondary antibodies (1:500, Thermo Fisher) were used for fluorescent detection of antibody labeling and Draq5 (1:5000, Biostatus) was used for nuclear staining. To obtain transverse views, stained embryos were manually sectioned using vibratome steel blades.

### Drug treatment

Embryos at the appropriate stage were treated in DMH1 (Sigma Millipore, D8696) at a final concentration of 10 μM in E3 fish water. Control embryos were treated similarly in an equal concentration of DMSO. Treated embryos were grown to the desired stage for analysis.

### Quantification of pSmad level

Embryos from an intercross of *hsp70:noggin3* and *nkx3.1:Gal4; UAS:NTR-mCherry* fish were heat shocked at 18 hpf, fixed at 24 hpf, and then stained with the pSmad antibody and the nuclear dye Draq5. *hsp70:noggin3* transgenic embryos were identified based on the absence of pSmad and mCherry expression. In wild-type embryos, the intensity of pSmad was measured within the Draq5^+^ nucleus of individual mCherry^+^ cells in both the dorsal and ventral sclerotome domains using the Fiji software (Schindelin et al., 2012). Similarly, in *hsp70:noggin3* embryos, the intensity of pSmad was measured in cells within the dorsal or ventral regions of the somite. Subsequently, the pSmad intensity in wild-type fish was normalized to that of *hsp70:noggin3* embryos and plotted on a graph.

### Statistical analysis

All the graphs were generated using GraphPad Prism software. Data were plotted as mean ± SEM. Statitical significance between two samples was determined using Mann-Whitney *U* test. *p* values: *p* > 0.05 (ns, not significant); *p* < 0.05 (*); *p* < 0.01 (**); *p* < 0.001 (***); *p* < 0.0001 (****).

## ACKNOWLEDGMENTS

We thank the zebrafish community for sharing reagents, particularly David Kimelman and Matthias Hammerschmidt. We are also grateful to Sarah Childs for sharing transgenic lines and offering critical input on this project, as well as to the members of the Childs and Huang laboratories for their valuable discussions. The Pax7 antibody developed by Atsushi Kawakami was obtained from the Developmental Studies Hybridoma Bank, created by the NICHD of the NIH and maintained at The University of Iowa, Department of Biology, Iowa City, IA 52242.

## COMPETING INTERESTS

The authors declare that no competing interests exist.

## FUNDING

This study was supported by grants to P.H. from the Canadian Institute of Health Research (PJT-169113), Canada Foundation for Innovation John R. Evans Leaders Fund (Project 32920), and Startup Fund from the Alberta Children’s Hospital Research Institute (ACHRI).

## SUPPLEMENTAL FIGURES

**Figure S1.**
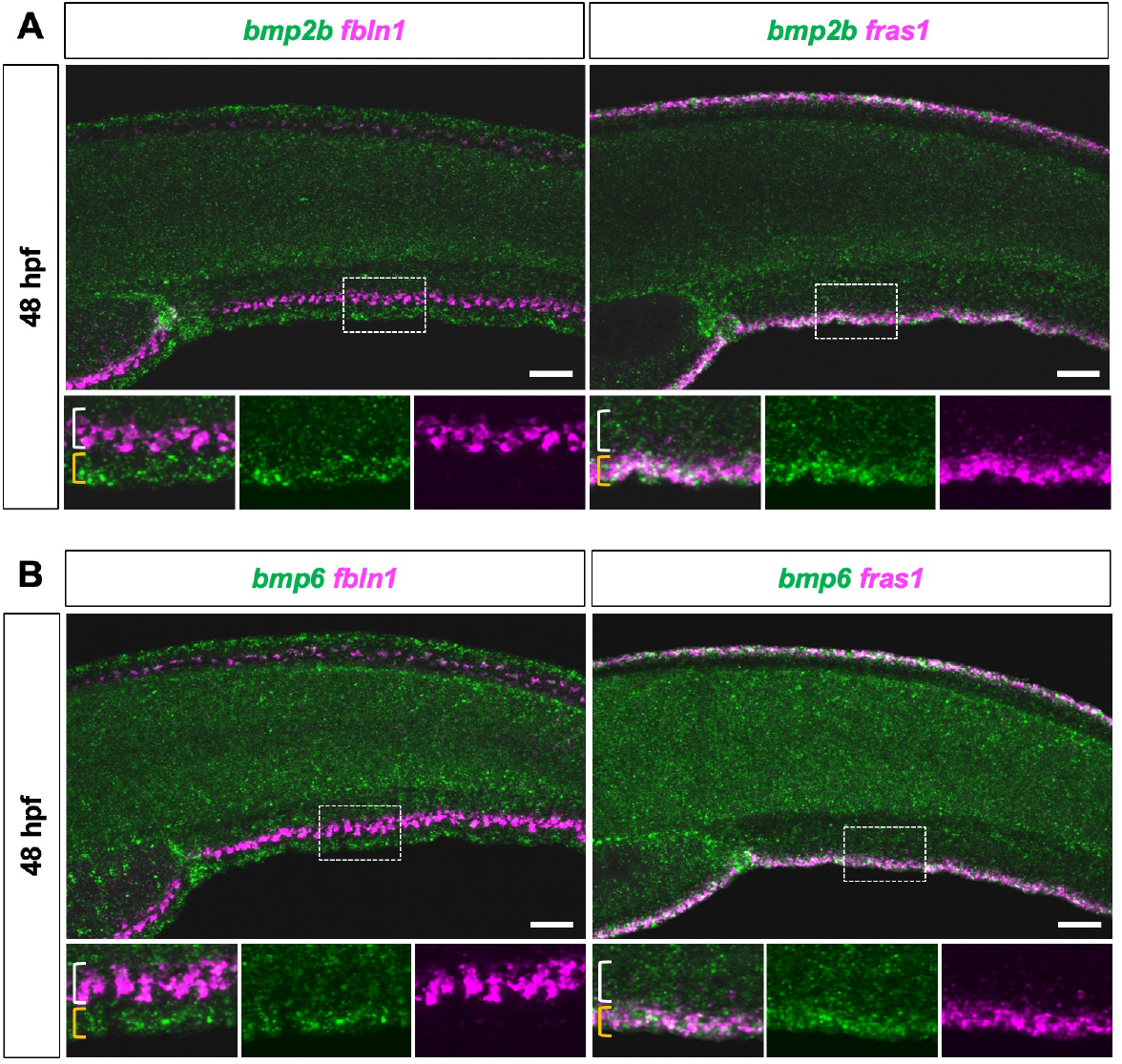
BMP ligands are expressed in the apical epidermal cells of the fin fold. (A-B) Double fluorescent in situ hybridization staining of *bmp2b* (A) or *bmp6* (B) (green) with *fbln1* or *fras1* (magenta) at 48 hpf. Both *bmp2b* and *bmp6* are expressed in *fras1^+^* apical epidermal cells (yellow brackets) adjacent to *fbln1^+^* fin mesenchymal cells (white brackets). Magnified views of boxed regions are shown in merged and individual channels. *n* = 15 embryos per staining. Scale bars: 50 μm.

**Figure S2.**
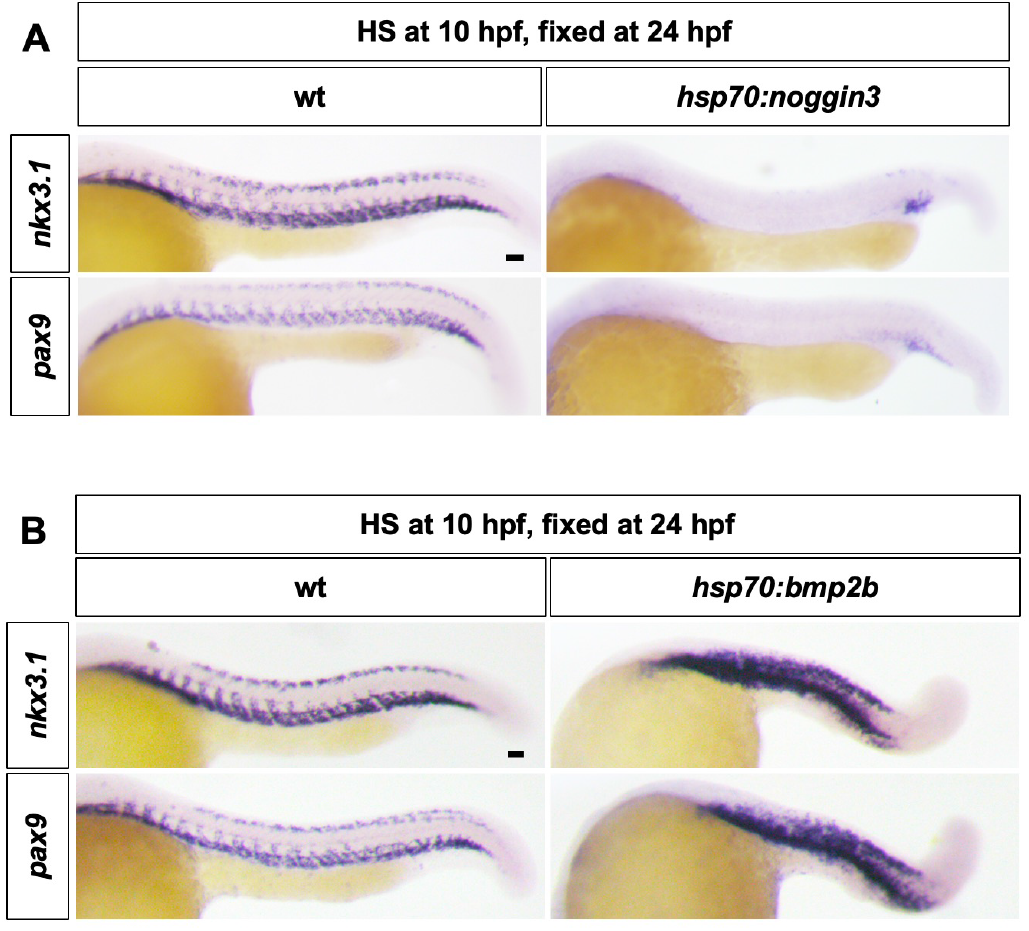
BMP signaling is necessary and sufficient for sclerotome induction. (A) Wild-type and *hsp70:noggin3* embryos were heat shocked at 10 hpf and stained at 24 hpf using *nkx3.1* and *pax9* probes. The expression of *nkx3.1* and *pax9* is completely absent in *hsp70:noggin3* fish. *n* = 15 embryos per condition. (B) Wild-type and *hsp70:bmp2b* embryos were heat shocked at 10 hpf and stained at 24 hpf for the expression of *nkx3.1* and *pax9*. The expression of *nkx3.1* and *pax9* is substantially expanded in *hsp70:bmp2b* fish. *n* = 15 embryos per condition. Scale bars: 50 μm.

## Notes

### Competing Interest Statement

The authors have declared no competing interest.

